# T-cell inflammation is prognostic of survival in patients with high-risk neuroblastoma enriched for an adrenergic signature

**DOI:** 10.1101/2023.06.26.546541

**Authors:** Maria E. Kaufman, Omar R. Vayani, Kelley Moore, Alexandre Chlenski, Tong Wu, Gepoliano Chavez, Sang Mee Lee, Ami V. Desai, Chuan He, Susan L. Cohn, Mark A. Applebaum

## Abstract

**Purpose:** T-cell inflammation (TCI) has been shown to be a prognostic marker in neuroblastoma, a tumor comprised of cells that can exist in two epigenetic states, adrenergic (ADRN) and mesenchymal (MES). We hypothesized that elucidating unique and overlapping aspects of these biologic features could serve as novel biomarkers.

**Patients and Methods:** We detected lineage-specific, single-stranded super-enhancers defining ADRN and MES specific genes. Publicly available neuroblastoma RNA-seq data from GSE49711 (Cohort 1) and TARGET (Cohort 2) were assigned MES, ADRN, and TCI scores. Tumors were characterized as MES (top 33%) or ADRN (bottom 33%), and TCI (top 67% TCI score) or non-inflamed (bottom 33% TCI score). Overall survival (OS) was assessed using the Kaplan-Meier method, and differences were assessed by the log-rank test.

**Results:** We identified 159 MES genes and 373 ADRN genes. TCI scores were correlated with MES scores (R=0.56, p<0.001 and R=0.38, p<0.001) and anticorrelated with *MYCN*-amplification (R=-0.29, p<0.001 and -0.18, p=0.03) in both cohorts. Among Cohort 1 patients with high-risk, ADRN tumors (n=59), those with TCI tumors (n=22) had superior OS to those with non-inflammed tumors (n=37) (p=0.01), though this comparison did not reach significance in Cohort 2. TCI status was not associated with survival in patients with high-risk MES tumors in either cohort.

**Conclusions:** High inflammation scores were correlated with improved survival in some high-risk patients with, ADRN but not MES neuroblastoma. These findings have implications for approaches to treating high-risk neuroblastoma.

## Introduction

Neuroblastoma is the most common extracranial pediatric malignancy and is hallmarked by heterogeneous patient outcomes. While low-risk tumors often spontaneously regress or differentiate, long-term survival rates for patients with high-risk neuroblastoma remain approximately 50% due to high rates of refractory and relapse disease (1). The current risk classification system within the Children’s Oncology Group utilizes several prognostic factors to guide treatment decisions including age at diagnosis, tumor histology, presence of *MYCN-*amplification, presence of 11q aberration, and DNA ploidy (2). However, with the exception of loss of 11q, none of these markers are prognostic within the high-risk group (3). Therefore, an improved knowledge of how tumor biology impacts response to therapy and ultimately survival is necessary.

High-risk neuroblastoma is comprised of two distinct cell lineages, mesenchymal (MES) and adrenergic (ADRN), defined by modifiable super-enhancers (SEs) that allow for lineage interconversion (4). Recent studies have noted that the MES lineage is associated with an immunogenic state (5,6). Studies in numerous cancers have shown that tumors with immunogenic microenvironments have better prognoses and are more responsive to immunotherapy (7). While neuroblastoma has been considered a “cold” cancer, recent studies have shown that the immune landscape in neuroblastoma is nuanced. *MYCN-*amplified tumors, which are comprised of a higher proportion of ADRN type cells, are typically associate with “cold” or low T-cell-inflamed (TCI) signatures and few tumor associated T-cells (8,9). Conversely, among patients with *MYCN*-non-amplified high-risk tumors, high TCI status is associated with improved survival (10).

Furthermore, MES cell lineage has also been associated with intra-tumoral TCI (5), and recent studies in cell lines have demonstrated that in response to inflammatory stimuli, MES cells release proinflammatory cytokines, leading to increased T-cell killing of tumor, while adrenergic cells do not (6). Overall, these findings suggest that the relationship between *MYCN*-amplification, TCI, and mesenchymal cell lineage is complex and interwoven and increased understanding of these relationships is vital to understanding the immune component of neuroblastoma.

In this study, we aimed to optimize the epigenetic characterization of neuroblastoma tumors as predominantly MES or ADRN by utilizing a novel approach combining ChIP-seq, KAS-seq, and RNA-seq to identify the most relevant lineage-specific super-enhancers. We then examined the association between TCI in MES versus ADRN tumors. We also evaluated the association between survival and TCI in tumors with different cell lineages to investigate the impact of inflammary cells in the tumor microenviroment in MES vs ADRN tumors.

## Methods

### Cell culture of neuroblastoma cell lines

ADRN neuroblastoma cell lines LA1-55n, SH-SY5Y, NBL-W-N, SK-N-BE2, and NBL-S and MES cell lines LA1-5s, SHEP, and NBL-W-S were grown at 5% CO_2_ in RPMI 1640 medium (Life Technologies) supplemented with 10% heat-inactivated FBS, 2 mmol/L l-glutamine, and 1% penicillin/streptomycin. Cells were counted using a hemocytometer and grown in a T225 flask for a period of 24 hours. Cells were then harvested and pelleted in 50 mL conical tubes. They were then washed with a 1X DPBS solution (Gibco 14190-144) and resuspended in 20 mL RPMI 1640 medium (Life Technologies). Resuspended cells were separated into 10 mL aliquots for chromatin extraction.

### Chromatin extraction and ChIP-seq

ChIP was performed as previously described^8^ with duplicate biologic replicates. Briefly, cultured cells were fixed with 36.5% Formaldehyde (Sigma F87750) to a final concentration of 1%. Nuclei were isolated and the DNA was sheared to 100-400 bp fragments. Histone-bound chromatin was immunoprecipitated using Anti-Histone H3K27ac antibody (Abcam ab4729) and H3K4me3 antibody (Cell Signaling 9751S). Cross-linking was reversed and DNA was purified using Qiagen PCR kit (Qiagen 28104). ChIP-seq libraries were made using Ovation® Ultralow System V2 kit (Tecan Genomics 0344NB-32) per manufacturer’s instructions and sequenced on an Illumina NovaSeq 6000.

### RNA-seq library construction

RNA was isolated with triplicate biologic replicates using TRIzol® reagent (Life Technologies) according to the manufacturer’s protocol. The concentration was measured using UV spectroscopy (DeNovix). DNA was removed with the TURBO DNA-free kit (Thermo Fisher Scientific) per the manufacturer’s instructions. Ribosomal RNA was removed with the oligo-DT kit, and a directional RNA library was constructed. 100 base-pair, paired-end libraries were sequenced on an Illumina NovaSeq 6000. Reads underwent quality control, trimmed as indicated, and aligned to hg38 using STAR RNAseq aligner (11).

### KAS-seq library construction

KAS-seq libraries were generated as previously described (12) in duplicate biologic replicates. Briefly, cells were incubated in completed culture medium containing N_3_-kethoxal. Cells were collected and genomic DNA (gDNA) was isolated from cells by PureLink genomic DNA mini kit (Thermo, K182002). gDNA was sheared and fragmented to 150-350 bp segments. 5% of the fragmented DNA was saved as input, and the remaining 95% was used to enrich biotin-tagged DNA. DNA was eluted and its corresponding input control were used for library construction by using Accel-NGS Methyl-seq DNA library kit (Swift, 30024). The libraries were sequenced on Illumina Nextseq500 platform with single-end 80-bp mode.

### Processing of ChIP-seq, KAS-seq, and RNA-seq sequencing

The quality for all sequencing files was verified using FASTQC v.011.5 (13) and cleaned using Trimmomatic v0.36 (14) using default settings. Reads were then aligned to GhC38 using Bowtie2 v2.3.0 (15) using default settings and deduplicated with PICARD v2.8.1 (16). Peaks were called using MACS2 v2.1.0 (17) for each sample using the arguments --keep-dup=auto --broad. For the RNA-seq, sequencing quality was verified using FASTQC v.011.5 and reads were cleaned using Trimmomatic v0.36. After alignment, lists of gene counts were generated for each cell line using featureCounts (18). Reads were loaded into R v3.6.0 using DESeq2 v1.20.0 (19) and genes differentially expressed between MES and ADRN cell lines were identified (p_adj_<0.05 and absolute log fold change >1). For analysis of gene expression, reads were processed using a variance stabilizing transformation with the DESeq2 package.

### Compiling super-enhancers using ROSE and identifying lineage-specific genes

Rank Ordering of Super-Enhancers (ROSE) (20) was performed for each cell line using the H3K27ac and H3K4me3 peaks with the arguments -c -s 0 -t 0. To identify super-enhancers, any H3K27ac peaks with signal >1000 ROSE units in the H3K4me3 data were discarded as they represented transcription start sites. Peaks around the *MYCN* amplicon were also discarded. The ROSE algorithm was re-run on the remaining H3K27ac peaks to generate a list of SEs for each cell line. These SEs were then processed through the KAS-seq pipeline (12) to determine which super-enhancers are single-stranded. All single-stranded super-enhancers (ssSEs) for the ADRN cell lines and the MES cell lines were then combined and overlapping sequences were merged within each phenotype to create list of ssSEs for each phenotype. Each ssSE only had to be identified in a single cell line to be retained. Genes within 500 kbs of the lineage-specific ssSEs were identified and retained if they also had significantly higher expression in the corresponding cell lineage. Retained genes were included as the ADRN and MES signatures, respectively. Pathway enrichment analysis was perfomed for the final signatures using Gene Set Enrichment Analysis (21).

### Data visualization

Matrices with the ROSE generated super-enhancer and single-stranded super-enhancer coordinates and were created for each sample. Heatmaps were then generated for each cell line using these matrices. Both matrix and heatmap creation were done using deepTools v3.5.1 (22).

### Primary tumor RNA-seq

Publicly available RNA-seq of primary neuroblastoma tumors was used in Cohort 1 (GSE49710 and GSE49711) and Cohort 2 (TARGET NBL, dbGaP: phs000467.v21.p8) cohort. Reads were downloaded from the NCBI Gene Expression Omnibus and converted to transcripts per million.

### Tumor scoring and characterization

To assess relative expression of cell lineage specific genes within each tumor, we used Gene Set Variation Analysis (GSVA), a tool which models pathway variation across samples in an unsupervised manner (23). Each sample was assigned both a MES and ADRN score using the respective gene signature based on the single-stranded super-enhancer-associated genes as described above.

An adjusted mesenchymal score (MES_adj_) was calculated by subtracting the ADRN score from the MES score. Each tumor was also assigned a TCI score based on a published CD8+ T-cell gene signature (24), which has been validated by immunohistochemistry and shown to correlate with response to immunotherapy using the same method.

### Survival analysis

Survival was assessed by the Kaplan-Meier method using the log-rank test. Differences were also determined using 3-year point estimates of overall survival (OS) and event-free survival (EFS) as well as 95% confidence interval (CI) were assessed. P-values <0.05 were considered significant.

## Results

### Single-stranded super-enhancers identify cell lineage specific genes

Using the H3K27ac and H3K4me3 data, ROSE identified 1,512 super-enhancers in MES and 1,951 super-enhancers in ADRN cell lines. Of the MES super-enhancers, 270 overlapped with KAS-seq peaks (**Figure 1A**). There were 500 genes with increased expression in MES compared to ADRN cell lines that were within 500kb of a SE. Of these genes, 159 were near SEs that were single-stranded and retained in the final MES signature (**Supplementary Table 1**). KAS-seq peaks were also identified corresponding to 512 single-stranded super-enhancers in the ADRN cells (**Figure 1B**). There were 746 genes with increased expression in ADRN compared to MES cell lines near a SE. Of these genes, 373 were near SEs that were single-stranded and retained in the final ADRN signature (**Supplementary Table 2**; **Figure 1C)**.

**Figure 1:**
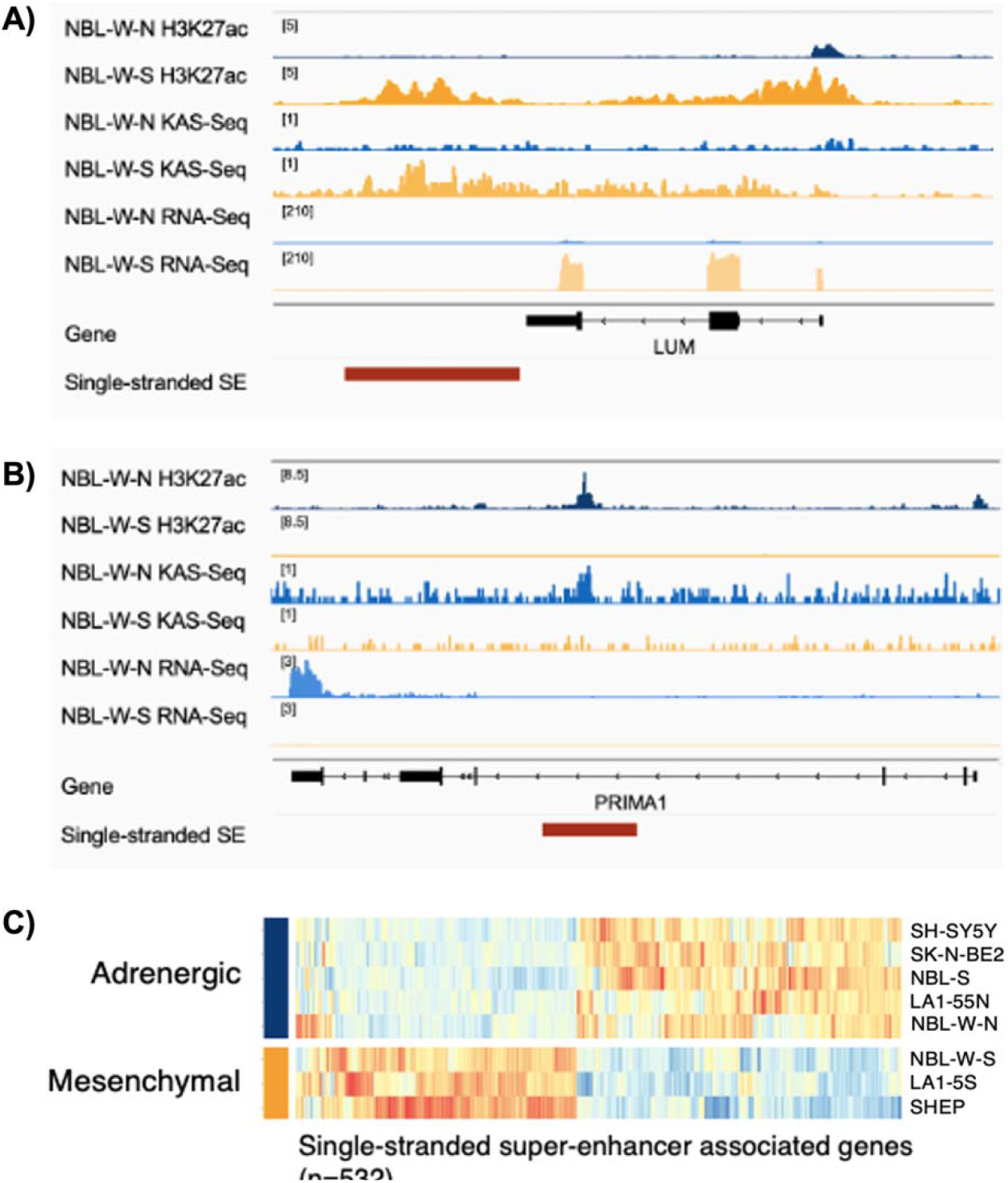
Single stranded super-enhancers identify highly expressed genes unique to the ADRN and MES lineages. A) H3K27ac deposition, KAS-seq pulldown, and RNA expression at MES single-stranded super-enhancer, denoted in red, associated with *LUM* in ADRN and MES isogenic cell lines. B) H3K27ac deposition, KAS-seq pulldown, and RNA expression at ADRN single-stranded super-enhancer, denoted in red, associated with *PRIMA1* in ADRN and MES isogenic cell lines. C) Heatmap representing 532 differentially expressed genes that were identified in five adrenergic (ADRN) and three mesenchymal (MES) cell lines in proximity to single-stranded super-enhancers (p<0.05). SE = super-enhancer.

We confirmed that MES and ADRN signature genes near ssSEs had higher expression compared to differentially expressed genes near a SE that was not single-stranded in 498 primary neuroblastoma tumors. In the 553 MES genes near a double-stranded SE, the average expression was 23.7 transcripts per million (TPM) versus 26 TPM for the 159 MES signature genes (p<0.001; **Figure 2A**). For the ADRN genes near a double-stranded SE, the average expression was 24.7 TPM versus 26.8 TPM in the 373 ADRN signature genes (p<0.001; **Figure 2B**). Similarly, in Cohort 2, the MES genes near a double-stranded SE had an average expression of 50.5 TPM versus 85.9 TPM in the MES signature genes (p<0.001). The ADRN genes near a double-stranded SE had an average expression of 69 TPM versus 87.8 TPM in the ADRN signature genes (p<0.001). Consistent with other published signatures (4), the genes in the ADRN signature were enriched for pathwyas related to the synaptic membrane (**Figure 2C, 2D**), while genes in the MES signature were enriched for pathways related to the extracellular matrix and collagen binding (**Figure 2E, 2F**).

**Figure 2.**
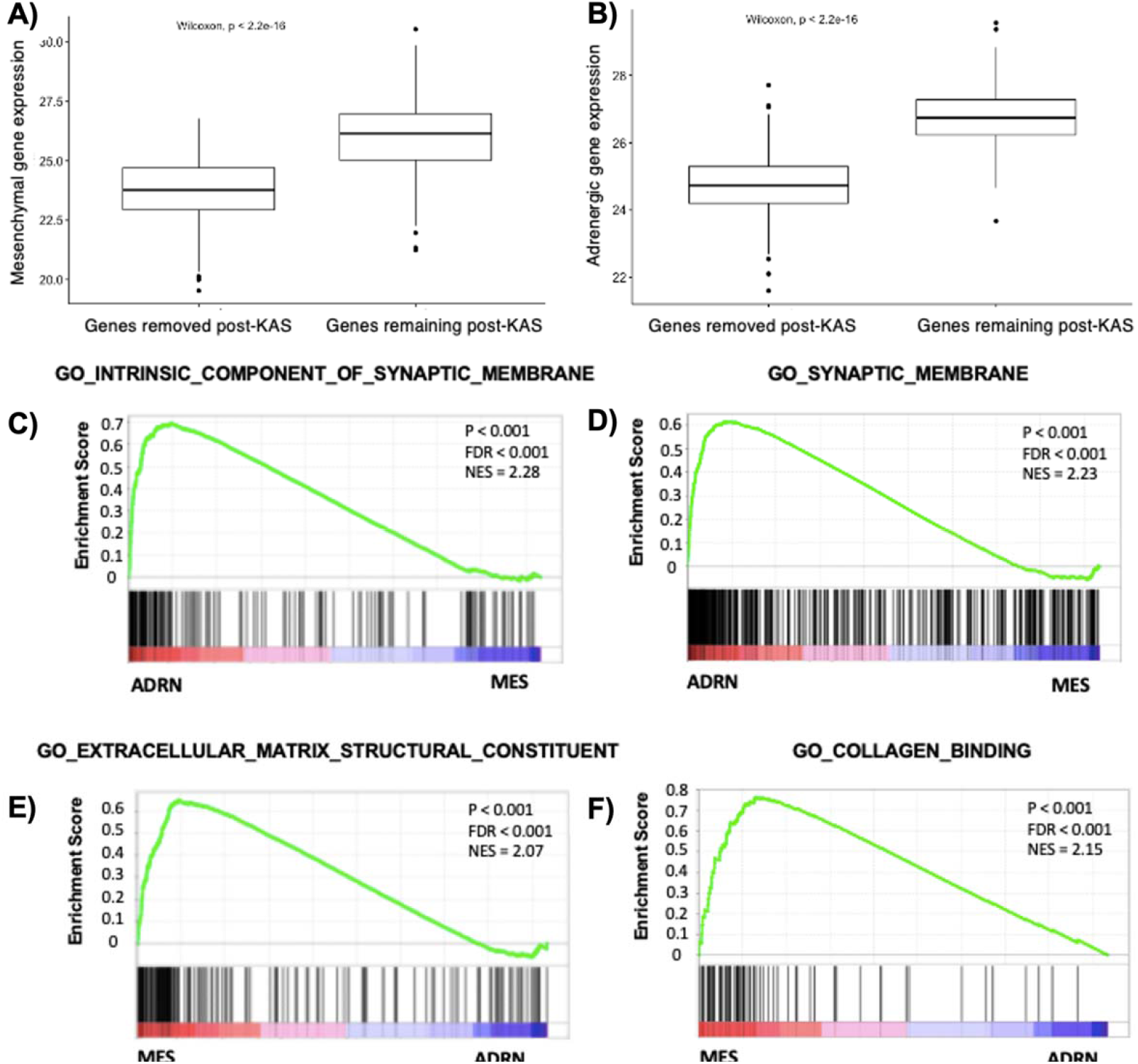
Lineage specific genes near ingle stranded super-enhancers have significantly higher expression and reflect tumor biology. A) Of the 712 genes with significantly higher expression in MES compared to ADRN cells near SEs, the 157 genes near ssSEs had higher expression in diagnostic biopsies than the 555 genes near double-stranded SEs. B) Of the 957 genes with significantly higher expression in MES compared to ADRN cells near SEs, the 371 genes near ssSEs had higher expression in diagnostic biopsies than the 586 genes near double-stranded SEs. C,D) ADRN signature genes were enriched for pathways related to their neural phenotype. (E, F) MES signature genes were enriched for pathways related to the extracellular matrix and collagen binding as has been described for these cell types.

### T-cell inflammation is correlated with MES cell lineage, *MYCN*-non-amplified tumor status, and low stage in primary neuroblastoma tumors

Using the final signatures derived from ssSEs, each tumor was assigned a MES score and an ADRN score using GSVA. To identify tumors with the strongest MES and lowest ADRN characteristics, we assigned an adjusted mesenchymal score (MES_adj_) generated by subtracting the ADRN score from the MES score. Among the 498 patients in Cohort 1, MES and ADRN scores were inversely correlated (R=-0.43, p<0.001; **Figure 3A, 3B**). However, while the expression profiles from the 145 tumors in Cohort 2 trended similarly, the finding was not statistically significant (R=-0.11, p=0.2). In both cohorts, TCI tumors had significantly higher MES_adj_ scores than NI tumors (p<0.001 and p<0.001, respectively; **Figure 3C, 3D)**.

**Figure 3.**
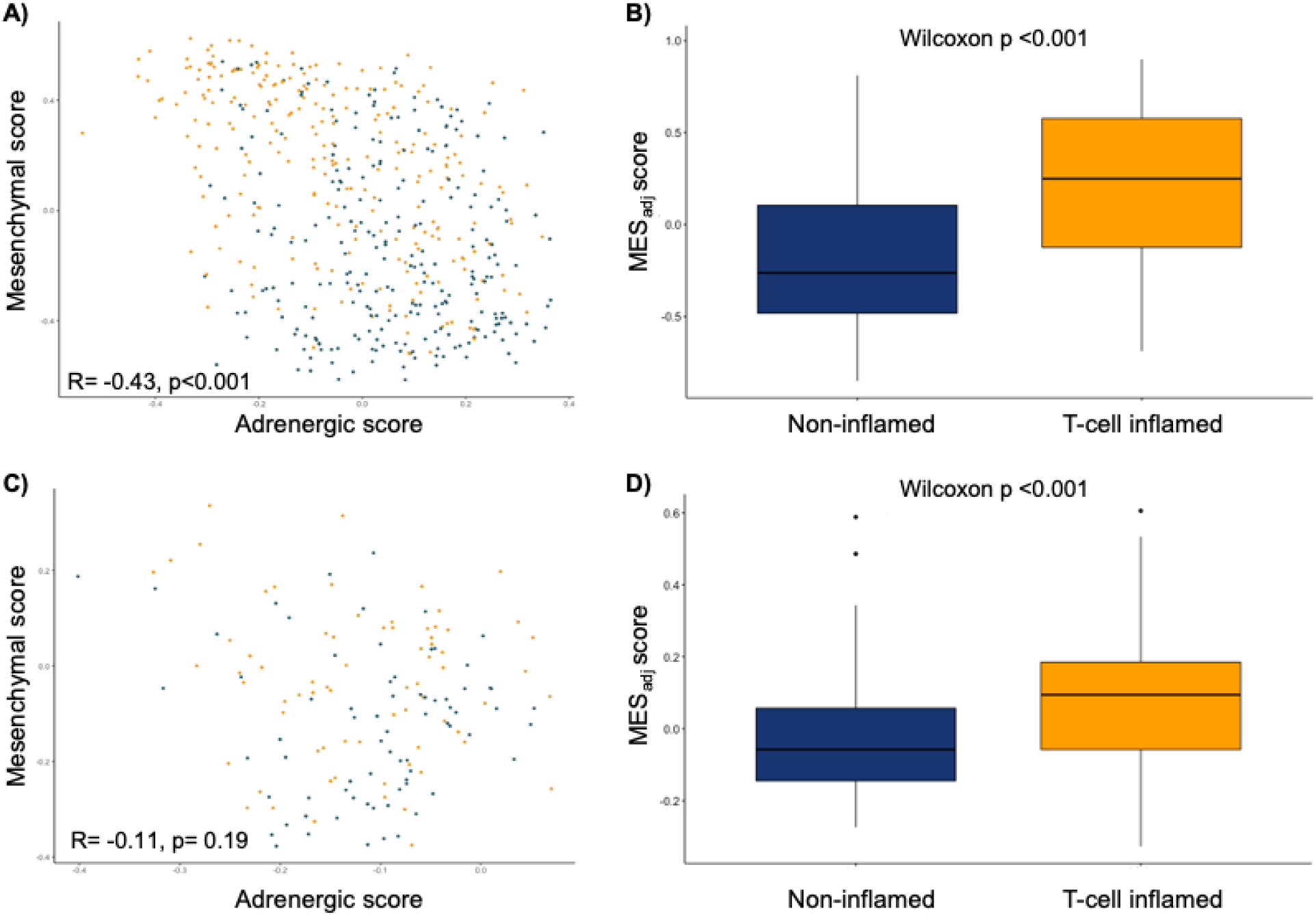
MES scores are anticorrelated with ADRN scores and correlated with T-cell inflammation (TCI). A) Association of MES and ADRN scores in Cohort 1. B) MES_adj_ scores were also significantly higher in TCI tumors in Cohort 1. C) Similar trends for inverse correlations were seen in Cohort 2. D) MES_adj_ scores were also significantly higher in TCI tumors in Cohort 2.

We also examined the relationship between TCI, MES_adj_, and other available clinical and biological features. In both cohorts, TCI was positively correlated with MES_adj_ (Spearman correlation=0.56, p<0.001 and 0.38, p<0.001, respectively). TCI was inversely correlated with *MYCN*-amplification in Cohort 1(biserial correlation=-0.29, p<0.001; **Figure 4A**) and Cohort 2 (biserial correlation=-0.18, p=0.03; **Figure 4B**). TCI and MES_adj_ were lower in high-risk tumors in Cohort 1 (p<0.001 and p=0.002, respectively), but these data were not available in the Cohort 2. TCI was also inversely correlated with higher International Neuroblastoma Staging System stage (ANOVA F=2.64, p=0.03, **Figure 4C**) in Cohort 1, although Stage 4S tumors also had low scores. Similarly, MES_adj_ was highest in stage 1 tumors and decreased by stage (p<0.001, **Figure 4D**), with and 4S also having high scores. Cohort 2 lacked low stage patients and no difference in TCI or MES_adj_ score was found between stage 3 and 4 tumors.

**Figure 4.**
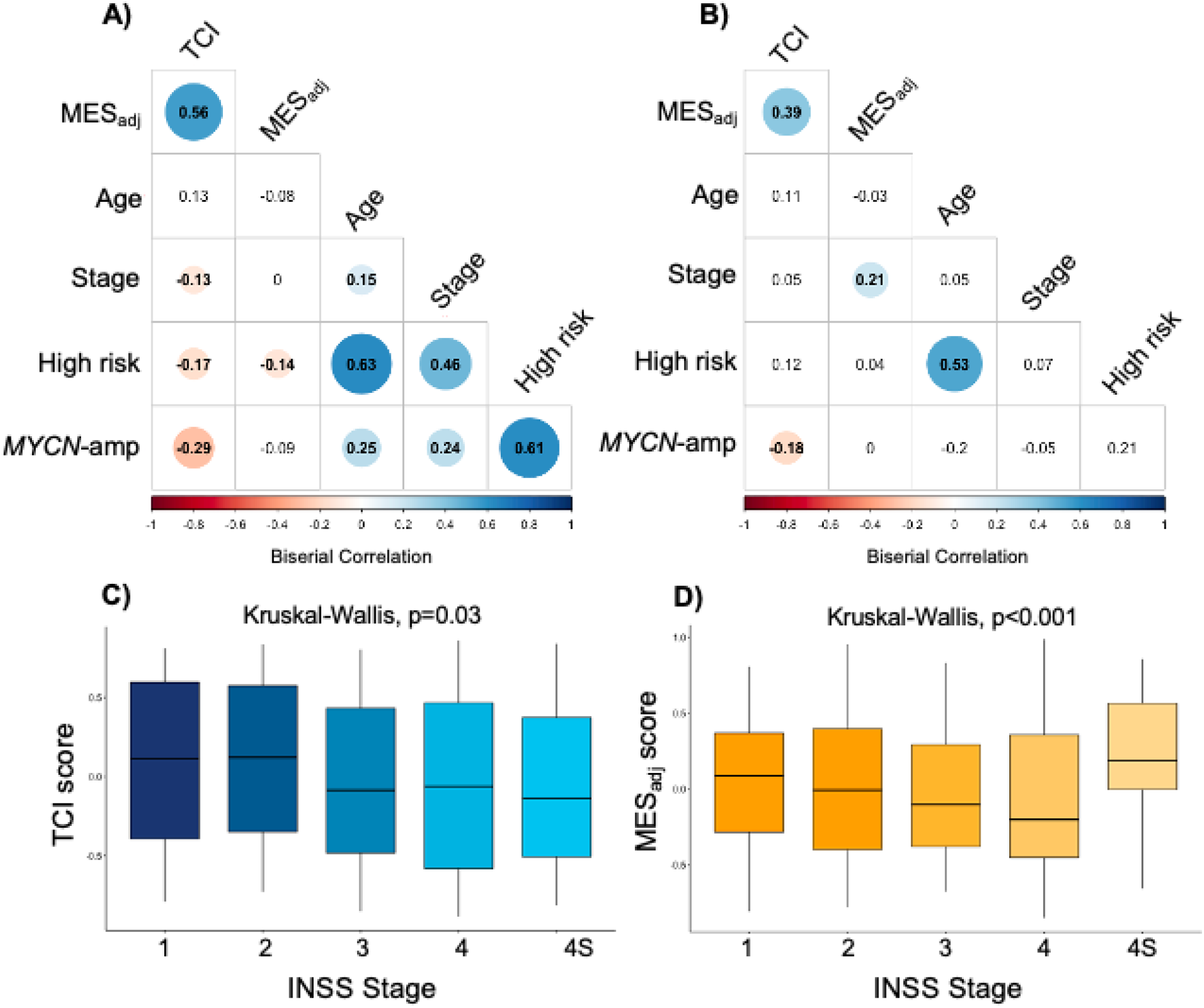
MES_adj_ and TCI scores are inversely correlated with features of high-risk neuroblastoma. MES_adj_ and TCI are correlated in both the A) Cohort 1 and B) Cohort 2 and inversely correlated with *MYCN*-amplificaiton, stage and age in Cohort 1. Colored circle representing the correlation coefficient shown only for those correlations with p<0.05. In Cohort 1, MES_adj_ and TCI decrease with increasing stage (C and D), although 4S tumors have higher MES_adj_ and lower TCI scores. Cohort 2 contained no stage 1 or 2 tumors and could not be compared.

### T-cell-inflammation is associated with improved survival in some high-risk patients with adrenergic tumors

To assess if tumors with the most distinct MES or ADRN-like scores were associated with patient outcomes, we next characterized tumors as MES (third with highest MES_adj_, n=59) or ADRN (third with lowest MES_adj_, n=59), and TCI (two thirds with highest TCI score, n=117) or NI (third with lowest TCI score, n=59) as described (8). As expected, the MES cohort was enriched for TCI (n=57), with just two patients classified as having NI tumors. Conversely, the ADRN cohort primarily of patients with NI tumors (n=37). This association was also observed in Cohort2, with the MES tumors (n=33) being mostly TCI (29/33 [87.9%]). Half of the ADRN tumors (16/32 [50%]) were TCI and half were NI (16/32 [50%]). In Cohort 1, the 22 high-risk patients with ADRN, TCI tumors had superior OS compared to 37 patients with ADRN, NI tumors (3-year OS 70%, 95% CI 53-93% vs 30%, 95% CI 19-50%; p=0.01; **Figure 5A**).

**Figure 5.**
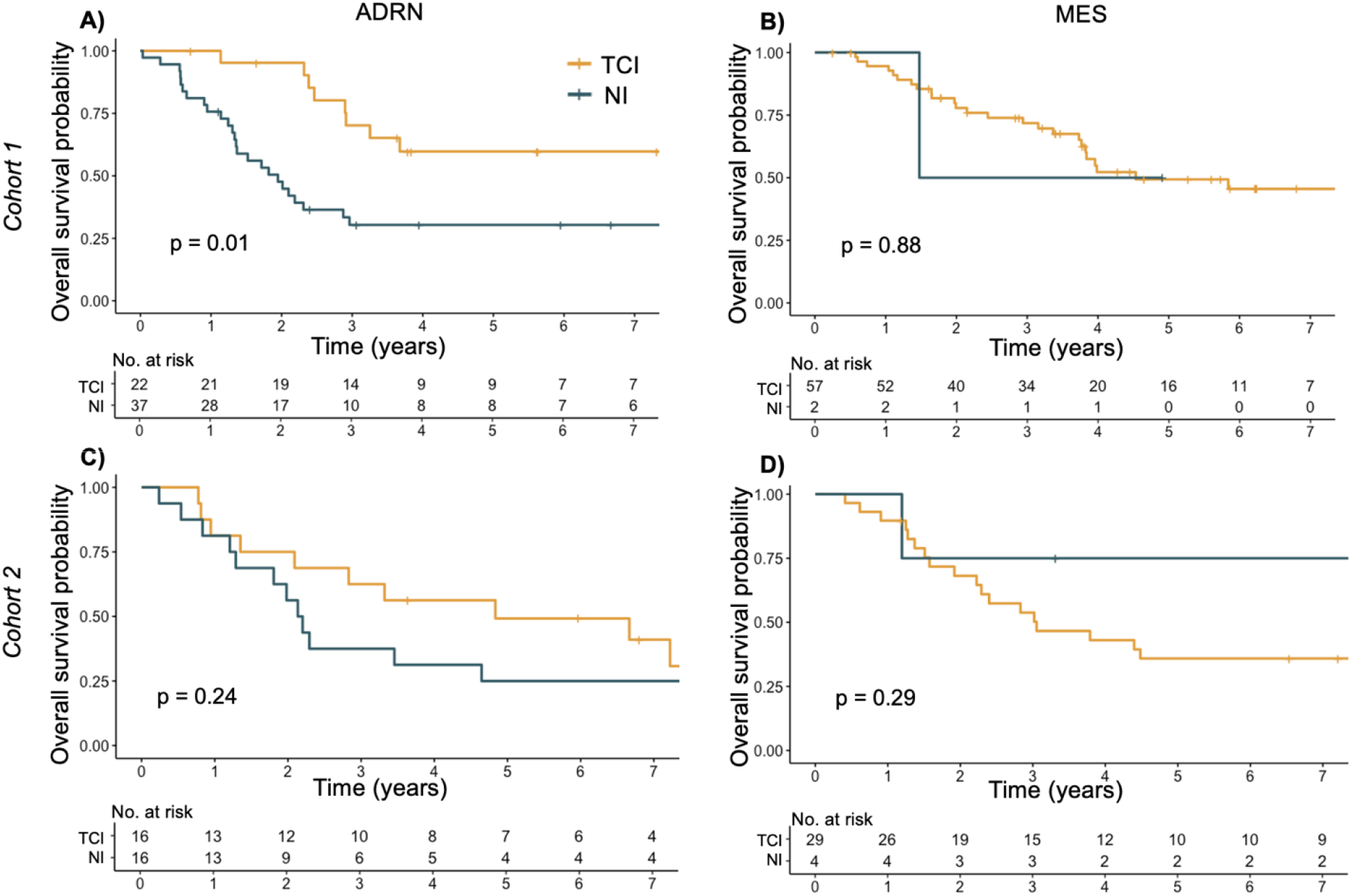
Inflammation score is prognostic for overall survival in high-risk, adrenergic tumors. A) Overall survival was significantly better for TCI tumors than NI tumors in patients woith ADRN tumors in Cohort 1 but not for B) those with MES tumors. C) Similar trends were seen in Cohort 2 for those with ADRN tumors, though these results were not statistically significant. D) In Cohort 2, there was no difference in overall survival according to inflammation status.

Conversely, OS among patients with MES tumors was not associated with inflammatory status (p=0.88; **Figure 5B**). In Cohort 2, while patients with high-risk, ADRN TCI (n=16) tumors possilby had a numerically better point estimate of OS than those with ADRN NI (n=16) tumors 3 years from diagnosis (3-year OS 63%, 95% CI 43-91% vs 38, 95% CI 20-70%; **Figure 5C**), the difference was not statistically significant, possibly due to small sample size. Patients with high-risk, MES tumors again had no difference in OS according to inflammation group (**Figure 5D**).

## Discussion

We generated novel lineage specific gene signatures in MES and ADRN neuroblastoma cell lines by incorporating transcriptionally engaged genes near single-stranded super-enhancers identified by KAS-seq analysis. By including genes with ssSEs, gene sets that were more highly expressed in cell lines and patient tumors were identified, highlighting the biologic relevance of this approach. We further show that the MES signature is strongly correlated with a T-cell inflamed signature whereas the ADRN signature is correlated with a non-immunogenic environment, though in a subset of tumors these associations were not observed. Finally, TCI signatures were correlated with outcome for patients with high-risk, ADRN tumors but were not prognostic among patients with MES tumors.

Biologic heterogeneity is a hallmark characteristic of neuroblastoma. Importantly, neuroblastoma cells are capable of interconverting between MES and ADRN with corresponding changes in phenotype and chemosensitivity (4,25,26). Furthermore, recent studies demonstrated that relapsed neuroblastoma tumors are enriched with MES cells, and several studies have focused on converting cellular phenotype to alter therapeutic sensitivity (27–29). While others have identified lineage specific gene signatures using lineage specific super enchancers (4,30), we also utilized KAS-seq data and incorporated genes with single stranded enhancers, a marker of active transcription, that were differentially expressed in the phenotypically distinct neuroblastoma cell types. This identified a unique set of genes more highly expressed than those identified only by analysis of classical histone marks, suggesting these genes are likely to play a more central role in neuroblastoma pathogenesis.

Improving our understanding of MES and ADRN phenotypes and their role in the pathophysiology of neuroblastoma tumors has been of intense interest (31). While neuronal and stromal cellular types have been recognized for decades (32), it has only been in the past few years that their relationship to patient tumors has come more into focus (27). Core regulatory circuitries for ADRN cells are well defined and include transcription factors such as ASCL1, DBH, ISL2, and DLK1, all of which were identified in our initial analysis, though ASCL1 was elimited during the KAS-seq pipeline (26,33). Beyond the key transcription factor PRRX1, lineage defining circuitries in MES cells have been harder to elucidate, suggesting a more complex network (30). Single cell sequencing experiments have verified that these two lineages do interconvert, but certain cells may be in a more committed ADRN path (34). Furthermore, conflicting evidence exists regarding the prognostic significance of signatures identifying the abundance of these cells in patient tumors. MES signatures in tumors at diagnosis have been shown to correlate with improved outcomes (27), but relapsed tumors and therapy resistant PDX models also have higher MES signature expression (35,36). In this study, we confirm that defining a precise relationship between ADRN and MES signatures in bulk sequencing and outcome is challenging, likely due to the relatively low abundance of MES cells in tumors due to their more senescent nature (37).

In both cohorts, we also identified an association between MES and TCI signatures. However, TCI signatures were associated with improved survival amongst patients with high-risk, ADRN tumors. This is consistent with a recent publication from Sengupta et al. where, using the same Cohort 1, they identified that tumors with higher MES scores were associated with activated immune states (5). They further demonstrated that MES cells promote TCI through cytokine secretion, are response to PD-L1 inhibition and are killed by cytotoxic T-cells. *MYCN*-amplification has been shown to be correlated with non-TCI tumors (8,38). In contrast to our findings, Bao et. al identified T-cell signatures that were predictive of outcome in neuroblastoma patients in a *MYCN* independent fashion.

In conclusion, we developed MES and ADRN signatures based on single-stranded super-enhancers which identified a set of genes highly expressed in multiple cohorts. We confirm the association of MES signatures and TCI and show correlations with improved outcomes for patients with TCI, ADRN signatures. Further studies are needed to determine if TCI tumors are more amenable to alterations in immunotherapeutic approaches.

## Data Availability Statement

ChIP-Seq, 5-hmC, KAS-Seq, and RNA-Seq raw and processed data for included neuroblastoma cell lines are available at GSE212882. For Cohort 1, RNA-Seq expression data and associated patient data was downloaded from GSE49710 and GSE62564 (SEQC-NB dataset). For Cohort 2, RNA-Seq raw read counts generated using HTSeq and clinical data from the TARGET-NBL project were downloaded from the GDC Data Portal (now available as STAR counts.)

## Author Contributions

Maria Kaufman: formal analysis, writing-original draft, writing-review and editing. Omar R. Vayani: formal analysis, writing-original draft, writing-review and editing. Kelley Moore: data curation and writing-review and editing. Alexandre Chlenski: data curation and writing-review and editing. Tong Wu: data curation and writing-review and editing. Gepoliano Chaves: writing-review and editing. Sang Mee Lee: data curation, analysis, and writing-review and editing . Ami V. Desai: writing-review and editing. Chuan He: writing-review and editing. Susan L. Cohn: writing-review and editing and supervision. Mark A. Applebaum: conceptualization, data curation, writing-original draft and writing-review and editing, and supervision.

